# Insights into the function of the chloroplastic ribosome-associated GTPase HflX in *Arabidopsis thaliana*

**DOI:** 10.1101/2023.03.03.530967

**Authors:** Marwa Mehrez, Cécile Lecampion, Hang Ke, Faten Gorsane, Ben Field

## Abstract

Ribosome-associated GTPases are conserved enzymes that participate in ribosome biogenesis and ribosome function. In bacteria, recent studies have identified HflX as a ribosome-associated GTPase that is involved in both ribosome biogenesis and recycling under stress conditions. Plants possess a chloroplastic HflX homolog, but its function remains unknown. Here, we characterised the role of HflX in the plant *Arabidopsis thaliana*. Our findings demonstrate that HflX does not have a detectable role in plant growth and development, nor does it play a distinct role in acclimation to several different stresses, including heat, manganese, cold, and salt stress. However, we found that HflX is required for plant resistance to chloroplast translational stress mediated by the antibiotic lincomycin. Our results suggest that HflX is a chloroplast ribosome-associated protein that may play a role in the surveillance of translation. These findings provide new insight into the function of HflX as a ribosome-associated GTPase in plants and highlight the importance of investigating conserved proteins in different organisms to gain a comprehensive understanding of their biological roles.

## Introduction

Chloroplasts are the organelles in plant and algal cells responsible for photosynthesis, the process that fuels plant growth and most life on earth by converting sunlight into chemical energy. Chloroplasts also host several other critical metabolic pathways, including de novo lipid biosynthesis, nitrogen and sulfur fixation, and hormone synthesis. In addition to playing important roles for their hosts, chloroplasts are also a major nutrient resource, containing a significant portion of the plant’s nitrogen and protein content, with Rubisco alone accounting for almost half of the soluble protein (Makino & Osmond, 1991; Eckardt et al., 1997).

Chloroplasts evolved from a symbiotic relationship between a cyanobacterium and a eukaryotic cell and have semi-autonomous features, including their own genome and gene expression machinery. The chloroplast translation machinery is well studied and strongly resembles that found in bacteria, with the addition of chloroplast specific proteins (Zoschke & Bock, 2018). In bacteria, a suite of ribosome associated and translational GTPases assist bacterial ribosome assembly, translation and ribosome turnover. Plant orthologues of these GTPases have been identified, although for many their molecular roles are poorly characterized (Suwastika et al., 2014; Mehrez et al., 2023). Certain are known to be essential, such as *SUPPRESSOR OF VARIEGATION11*, a plant homologue of the translation GTPase elongation factor TU (EF-TU) (Liu et al., 2019), and ObgC, a plant homologue of the bacterial ribosome-associated GTPase Obg(Chigri et al., 2009; Bang et al., 2012). The ribosome-associated GTPase high frequency of lysogenization X (HflX) has recently received attention in both prokaryotes and animals. HflX binds to 50S ribosomal subunits (Jain et al., 2009), is implicated in ribosome biogenesis (Schaefer et al., 2006) and also acts as a ribosome splitting factor (Zhang et al., 2015; Coatham et al., 2016; Dey et al., 2018; Rudra et al., 2020). While not essential for growth under standard conditions in *E. coli*, HflX is required for acclimation to heat stress where its ribosome splitting activity allows recycling of stalled ribosomes (Zhang et al., 2015; Dey et al., 2018). Interestingly, in addition to its GTPase activity, HflX may also use an ATP-dependent RNA helicase activity for the repair and reactivation of heat-damaged ribosomal RNA (Dey et al., 2018). Animals possess an HflX ortholog known as GTPBP6. Like HflX, GTPBP6 is a ribosome recycling factor, and is required for the assembly of mitochondrial ribosomes (Lavdovskaia et al., 2020; Hillen et al., 2021). In contrast to animals and bacteria, no role has been attributed to the plant HflX. However, the enzyme is reported to show a chloroplastic localization (Suwastika et al., 2014) and is found in association with the 50S subunit of the chloroplast ribosome (Olinares et al., 2010).

In this study, we focus on the physiological function of the plant HflX. We show that the canonical HflX shares strong structural conservation with bacterial HflX enzymes. We also report a second non-canonical plant HflX-like enzyme that has independent evolutionary origins. Using independent T-DNA insertion mutants, we show that the canonical HflX is dispensable for normal growth and development. Although HflX seems not to be involved in acclimation to a range of stress conditions, we find that it is required for resistance to lincomycin, an antibiotic that inhibits chloroplast translation. On the basis of this, we suggest that HflX is able to protect translation machinery either via blocking lincomycin binding to the ribosome, or promoting the recycling of ribosomes stalled by lincomycin. Altogether, our results suggest that HflX is a chloroplast ribosome-associated enzyme that plays a role in chloroplast translation, while its precise contribution to plant growth and stress acclimation remains uncertain.

## Material and Methods

### Plant material and growth conditions

The wild type was Col-0. SALK_002001C (*hflx1-1*), SALKseq-041831.1 (*hflx1-2*) and SALK-057030.1 (*hflx1-3*) were provided by the Signal Insertion Mutant Library (http://signal.salk.edu) (Alonso et al., 2003). Homozygous insertion mutants were isolated and confirmed by PCR. For growth in normal conditions, seeds were sown in soil and transferred to separate pots 7 days after germination. Plants were grown either under long day conditions (16h light/ 8h dark) or short day conditions (8h light/ 16h darkness), at 22/18°C with 120 μmol photons m^−2^ s^−1^ lighting. For growth in culture dishes, seeds were surface sterilized in 70% ethanol containing 1% sodium hypochlorite and 0.005% Tween 20 for 10 min, then washed with 100% ethanol, dried and transferred in a grid pattern onto square plates containing 50 ml of MS/2 medium (0.5× Murashige and Skoog salts [Merck Sigma-Aldrich], 1% sucrose, 0.5 g/L MES, and 0.8% agar, adjusted to pH 5.7 with KOH). After 2 days of stratification at 4°C, plates were placed in a culture room with 16h light (at 22°C)/8h darkness (at 19.5°C) and 80 μmol photons m^−2^ s^−1^ lighting.

### Stress treatments

We used the growth conditions mentioned above unless otherwise indicated. For heat stress, seedlings were grown on MS/2 for 12 days in standard conditions. Plates were transferred in a Percival and exposed to heat treatment for 24h at 40°C, then, allowed to recover in standard conditions.

For manganese stress, seedlings were grown on MS/2 (with 0.4% phytagel (Sigma-Aldrich) instead of 0.8% agar) supplemented or not with 2mM filter sterilized MnSO_4_.

For cold stress, seeds were grown on MS/2 for 7 days in standard conditions with 50 μmol photons m^−2^ s^−1^ lighting. Plates were then either kept in standard conditions (control) or transferred to a cold room at 5°C with 40 μmol photons m^−2^ s^−1^ lighting.

For salt stress, seedlings were sown on MS/2 without sucrose and grown for 7 days and then transferred to MS/2 without sucrose supplemented or not with 150 mM NaCl.

For lincomycin treatment, seeds were germinated on MS/2 containing 35 μM filter sterilized lincomycin.

### Genotyping

DNA extraction was based on the approach of Edwards et al. (1991) with modifications. DNA from one leaf disk was extracted in 400 μl of DNA extraction buffer (200 mM Tris-HCl, 250 mM NaCl, 25 mM EDTA, 0.5% w/v SDS, 20 μg/ml RNAse, pH 7.5) using a pellet pestle in 1.5ml tube. The samples were incubated for 1 h at 65°C and centrifuged at 3000 g for 10 min. The supernatant was transferred to a new tube containing 200 μl of phenol-chloroform-isoamyl alcohol (25: 24: 1). Tubes were inverted several times, left for 5 min and centrifuged at 3000 g for 10 min at 4°C. the upper aqueous phase was transferred to a new tube to which an equal volume of isopropanol was added. The tubes were then left for 1 h at room temperature. After centrifugation at 4000 g for 25 min, the supernatant was discarded and the pellet was washed with 70% ethanol. The ethanol was completely removed, and the pellet was air dried and resuspended in TE buffer (10 mM Tris-HCl, 1 mM EDTA, pH 8.0). T-DNA insertions were then analysed using specific primers (Table S1) in PCR reactions with Emerald Master Mix (Takara). PCR conditions were as follows: an initial step at 98°C for 30 s, followed by 38 cycles of 98°C for 15 s, 58°C for 20 s and 72°C for 1 m-1 m 30 s.

### RNA extraction, RT-PCR and qRT-PCR

RNA extraction was performed using Tri-Reagent (Sigma-Aldrich) and quality confirmed by agarose gel-electrophoresis. RNA was treated with DNAse I (Thermo scientific), and cDNA was synthesized from 500 ng of RNA using Primescript RT Reagent Kit (Takara) with random hexamer primers. RT-PCR was performed as mentioned above for genotyping reactions using specific primers (Table S1). qRT-PCR was carried out in a Bio-Rad CFX96 real-time system using the following conditions: 95°C for 30 s, followed by 44 cycles of 95°C for 5 s, 59°C for 30 s and 72°C for 30 s. Each qRT-PCR reaction was performed in 15 μl reaction volume that consisted of 1 μl of cDNA [12.5 ng/μl], 2.4 μl of primer mixture (2.5 μM for each primer) (Table S1), 7.5 μl TB Green Premix Ex Taq II (Tli RNaseH Plus) (Takara). Melting curves were performed to confirm amplification specificity.

### Plant growth measurements

For rosette area measurements, plants grown on soil were photographed at different times during their growth using a camera (Panasonic Lumix, 20-1200). Images were then automatically analysed using the ARADEEPOPSIS pipeline (Hüther et al., 2020). For measurements of seedling area, plates were scanned at the indicated time. The images were then analysed in ImageJ (NIH) and the area from each seedling was obtained.

### fluorescence analysis

Chlorophyll fluorescence was measured in a Fluorcam FC 800-O imaging fluorometer (Photon System Instruments). The plants were adapted to dark for 20 min and the PSII maximum quantum yield (Fv/Fm) was calculated as (Fm − Fo)/Fm.

### Chlorophyll quantification

Chlorophyll quantification was performed as previously described (Sugliani et al., 2016). Briefly, chlorophyll was extracted from frozen seedlings homogenised using pellet pestle in ice cold 90% acetone saturated with sodium carbonate and kept overnight at -20°C. When the plant material was completely white, the samples were centrifuged and the supernatant was transferred to a new tube. The absorbance was measured between 350-750 nm using an 80% acetone blank in a Varian Cary 300 spectrophotometer (Agilent). Total chlorophyll content was calculated using a full spectra fitting algorithm (Chazaux et al., 2022). For each line, 5 biological samples from different plates were used for the absorbance measurements. The experiment was repeated twice.

### Phylogenetic inference and protein structure analysis

Using *E. coli* HflX as a query, homologous proteins from photosynthetic organisms were identified by BLAST search using public data at the National Center for Biotechnology Information (NCBI) and JGI. The representative bacterial HflX/HflXr and animal homologs were previously identified (Suwastika et al., 2014; Koller et al., 2022). Multiple-sequence alignments were performed using MAFFT v7.40262 with option –auto (Katoh et al., 2019). Phylogenetic reconstructions were created using maximum likelihood with the IQ-TREE web server version 1.6.12 using default settings, with LG +F+ I + G4 automatically selected as the best fit evolutionary model based on BIC values by ModelFinder (Trifinopoulos et al., 2016). Branch support was tested using two methods: ultrafast bootstrap approximation using 1000 bootstraps, and the non-parametric Shimodaira–Hasegawa–like approximate likelihood-ratio test (aLRT). The alignments and trees are available in supplementary file 1.

For structural analysis protein structures were retrieved from RSCB Protein Data Bank (PDB) or the EMBL-EBI Alphafold database(Jumper et al., 2021; Varadi et al., 2022) and visualised using ChimeraX (Pettersen et al., 2021).

### Data analysis

Data analysis and visualisation was conducted in R using scripts previously described (Romand et al., 2022) with minor modifications. Data generated from ARADEEPOPSIS was analysed using the script provided in https://github.com/cecile-lecampion/Analyse_croissance. For each experiment, plates were considered as independent replicates and individual plants were considered as biological replicates. The experiments were performed at least twice and similar results were obtained. The significance of differences in categorical data (cotyledon death) was calculated using the proportion test as previously described (Romand et al., 2022).

## Results

### *Arabidopsis thaliana* contains two HflX homologs with different evolutionary origins

We analysed the distribution of HflX enzymes in plants, bacteria and other organisms. We found that green plants possess two HflX homologs (Fig. 1a). Phylogenetic analysis confirmed that the previously identified HflX groups with the green non-sulfur bacteria rather than cyanobacteria (Suwastika et al., 2014). This plant and algal HflX clade includes an HflX from the unicellular red alga *Cyanidioschyzon merolae*, strongly suggesting an evolutionary origin close to the emergence of chloroplast-containing organisms. Surprisingly, we found a second group of plant HflX-like enzymes that forms a separate clade with distinct evolutionary origins. Interestingly, this plant HflX-like clade appears to be closely related to the animal HflX enzymes, as well as an HflX from the thermophilic archaeon *Sulfolobus solfataricus*. Interestingly, we note that the plant HflX-like enzymes are not canonical because they lack the conserved C-terminal domain (CTD) found in animal and bacterial HflX enzymes (Fig. S1).

**Figure 1.**
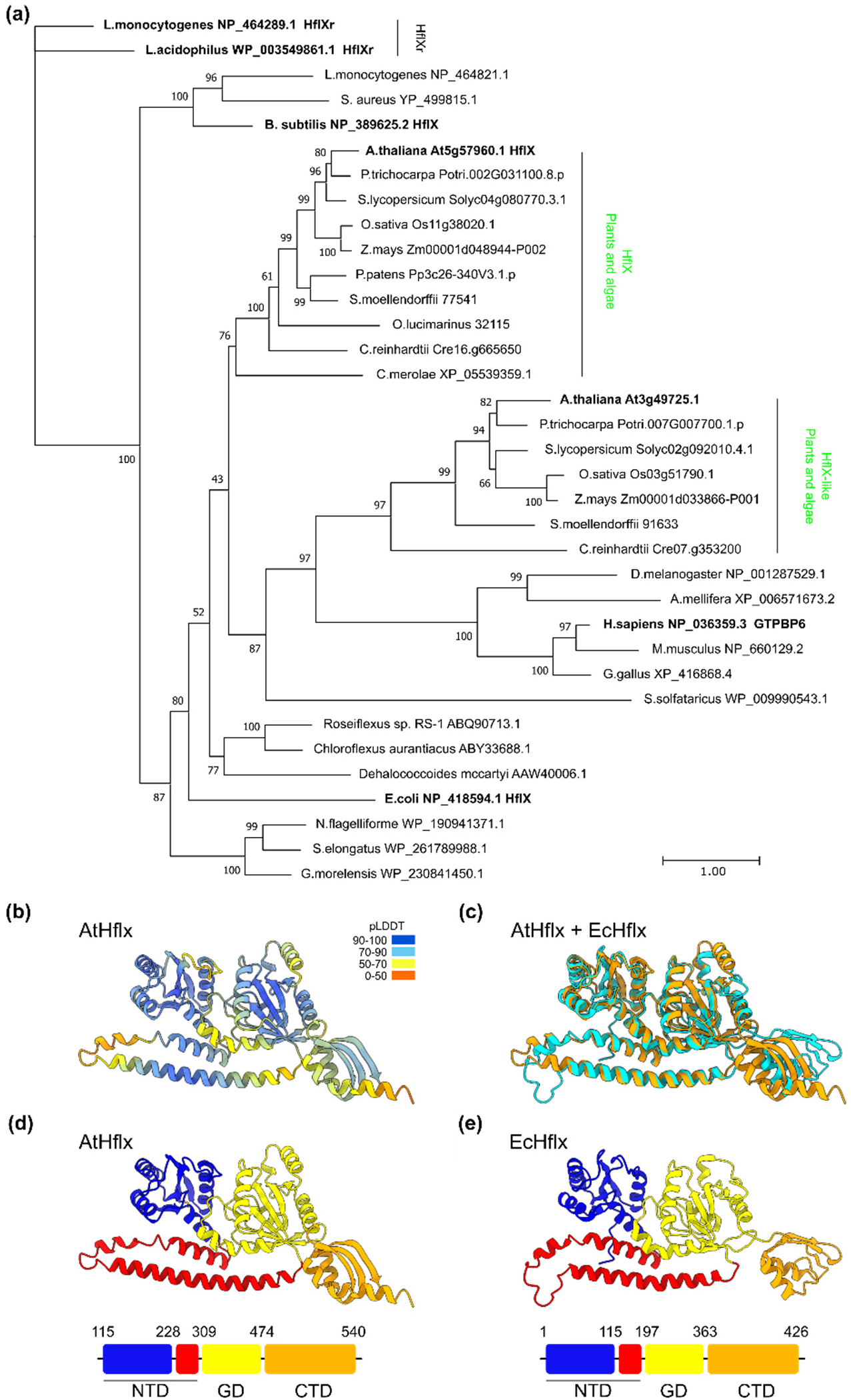
Arabidopsis contains an HflX homolog with a conserved structure. (a) A maximum-likelihood phylogenetic tree of selected HflX proteins from eukaryotes and prokaryotes. Plant and algal HflX clades are indicated. The scale bar indicates substitutions per site, and statistical support for branches is shown at the nodes. (b) Alphafold model of Arabidopsis HflX (AtHflX, Q9FJM0) with confidence-per-residue coloring (pLDDT). The predicted chloroplast transit peptide is not shown. (c) Arabidopsis HflX model (orange) aligned with the structure of ribosome-associated *E. coli* HflX (cyan)(PDB 5ADY)(Zhang et al., 2015). Domain organisation of the (d) Arabidopsis and (e) *E. coli* HflX enzymes. NTD, N-terminal domain; GD, G-domain; CTD, C-terminal domain.

Recently, HflXr enzymes required for enhanced antibiotic resistance were discovered in *Listeria monocytogenes* bacteria (Duval et al., 2018; Koller et al., 2022). Our phylogenetic analysis shows that plants and algae clearly lack HflXr orthologues.

Next, we analyzed the Alphafold predicted protein structure of the canonical Arabidopsis HflX (Fig. 1b). Arabidopsis HflX displays the same domain organisation as the *E. coli* HflX with the presence of a conserved NTD (N-terminal domain), GD (G domain) and CTD (C-terminal domain) (Fig. 1d & e). The NTD and GD showed strong similarities at the structural level when compared with the ribosome-bound *E. coli* HflX, with the exception of a small alpha-helix extension in the GD (Fig. 1c). The NTD of *E. coli* HflX interacts with the 23S rRNA on the 50S subunit. It is therefore likely that Arabidopsis HflX is able to interact with the 50S ribosomal subunit in a similar fashion to the *E. coli* HflX. The CTD showed more differences, with an altered orientation and an additional alpha-helix. Residues considered important for ATPase (corresponding to *E*.*coli* HflX Arg90 and Asp102) and GTPase activity (Gly252 and Ser343) are also conserved (Lavdovskaia et al., 2020) suggesting that the Arabidopsis HflX may be able to perform both ribosome splitting and helicase functions. Both the sequence and structural conservation suggest that the canonical Arabidopsis HflX is therefore able to play a similar role to bacterial HflX enzymes.

Despite belonging to the HflX family, the plant HflX-like enzymes show major differences with respect to canonical HflX enzymes (Fig. S1). Arabidopsis HflX-like possesses a conserved NTD and GD core, yet lacks a C-terminal region resembling the HflX CTD. This is instead replaced with an unstructured tail. In addition, there is an enlarged loop (N-loop) in the NTD. In *E*.*coli* HflX, the N-loop extends into the peptidyl-transferase center (Zhang et al., 2015). The enlarged N-loop of the HflX-like NTD is therefore likely to profoundly alter the manner in which the enzyme is able to interact with ribosomes. Finally, the localisation of HflX-like is not yet resolved. Indeed, Target P predicts a chloroplastic localisation (likelihood=0.63) (Almagro Armenteros et al., 2019), while the protein itself was identified in mitochondrial ribosome fractions (Rugen et al., 2019).

### The canonical HflX is not required for growth under standard conditions

To identify the role of the canonical *HFLX* in plants, we analyzed three *HFLX* T-DNA insertion mutants that we named *hflx 1-1, hflx 1-2* and *hflx 1-3* (Fig. 2a). We isolated homozygous lines (Fig. 2b), and confirmed the absence of a full-length *HFLX* transcript in the three mutants (Fig. 2c). This indicates that the three mutants are unable to produce the full length protein, and are likely to be loss-of-function knockouts.

**Figure 2.**
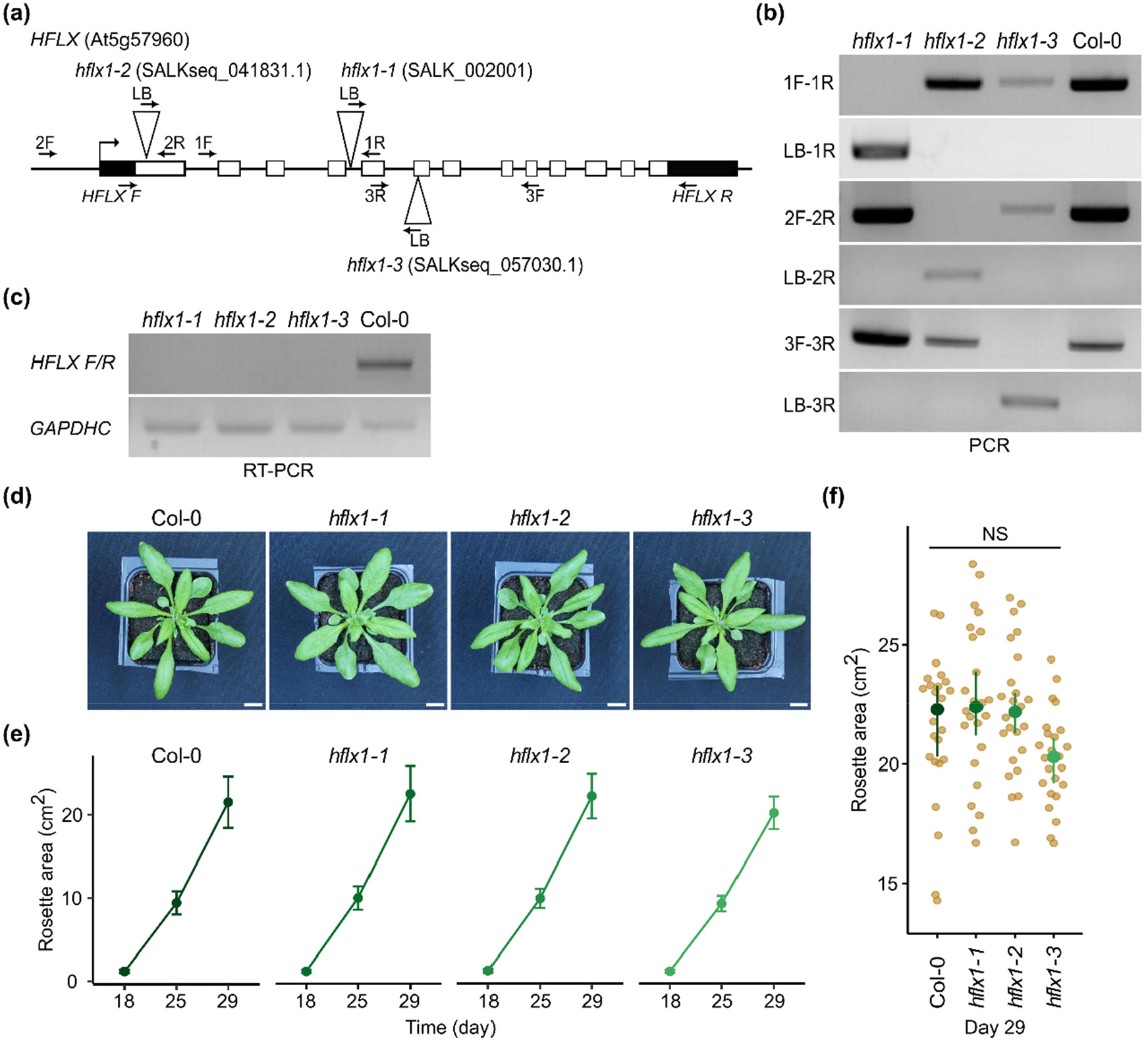
HflX is not required for normal growth. (a) Localization of T-DNA insertions in the canonical *HFLX* gene. Arrows indicate the position of genotyping primers. (b) Genotyping of *hflx* mutants using primers shown in (a). (c) RT-PCR amplification of the full-length *HflX* cDNA in the wild type Col-0 and the three *HflX* insertion mutants. (d-f) The phenotype of wild type (Col-0) and *hflx* mutants grown in long day conditions. (d) Photographs of plant rosettes at day 29, (e) quantification of vegetative growth rates, and (f) comparison of rosette area at day 29 (n= 24 plants per genotype). Scale bar, 1cm. Graphs show mean and 95% CI. NS, not significant.

To examine whether *HFLX* is required for plant growth, *hflx* mutants and the wild type Col-0 were grown in standard conditions and the growth rate was quantified. The three mutants showed no significant different in rosette size from the wild type under long day conditions (Fig. 2d-f). The results were similar in short day conditions, with the exception of *hflx 1-3* which was significantly smaller than all the other lines at day 39 (Fig. S2). As the *hflx 1-3* phenotype was not observed in the other mutants it cannot be explained by the knockout of *HFLX*. We conclude that HflX does not detectably influence the normal growth and development of Arabidopsis.

### *hflx* mutants are not hypersensitive to abiotic stress

In *E. coli*, the *hflx* mutant is hypersensitive to heat shock (Zhang et al., 2015) and HflX exhibits an ATP-dependent helicase activity that is necessary for RNA unwinding and rescuing heat-damaged 50S subunits (Dey et al., 2018). The structural similarity between *E. coli* and Arabidopsis HflX might suggest conservation of a role in acclimation to heat-shock. Therefore, we investigated the resistance of the three *hflx* mutants to a heat shock (HS) treatment at 40°C for 24 hr. One day after the HS, *hflx* mutants and wild type plants became pale green and showed evidence of cotyledon death (Fig. 3a). The efficiency of photosystem II (Fv/Fm) also decreased in response to HS, although there was no difference between the *hflx* mutants and the wild type (Fig. 3b). These results revealed that Arabidopsis HflX is not required for acclimation to HS.

**Figure 3.**
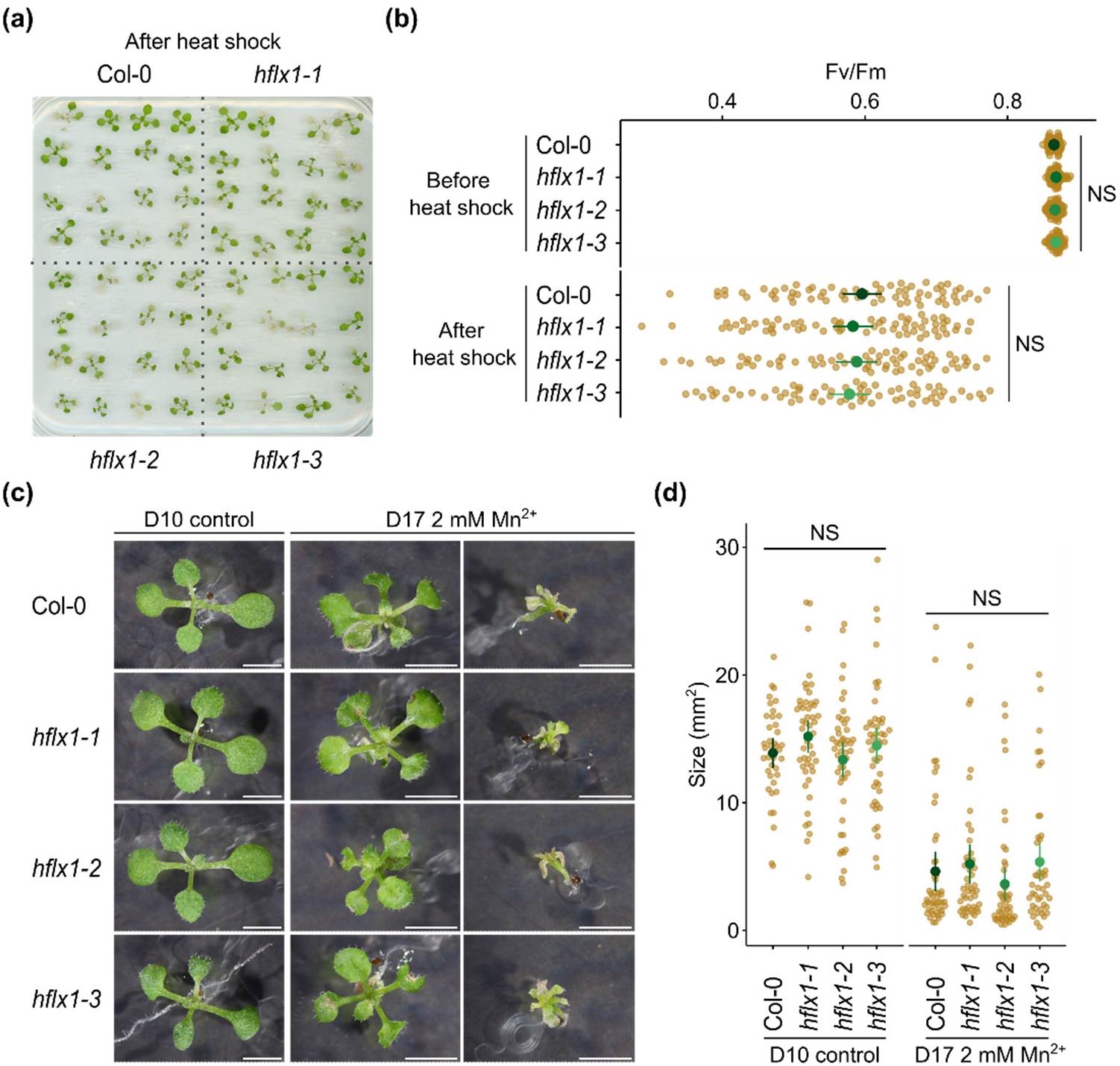
HflX is not required for acclimation to heat or excess manganese. (a) 12-day-old seedlings were subjected to a heat shock treatment for 24h at 40°C and photographed after one day of recovery. (b) PSII maximal efficiency (Fv/Fm) was measured in seedlings before and after heat shock treatment, n=64 plants per genotype. (c) Seedlings were grown on medium supplemented or not with manganese and photographed at the indicated day. Scale bar, 3mm. (d) Comparison of plant size of seedlings grown on medium with or without manganese at the indicated day. n= 44-47 plants per genotype. Graphs show mean and 95% CI. NS, not significant.

In *E. coli, hflx* mutants are also hypersensitive to excess manganese, a stress characterized by growth arrest, filamentation and lower rates of replication (Kaur et al., 2014; Sengupta et al., 2018). Therefore, we investigated the effect of excessive manganese on the growth of *hflx* mutants by germinating seeds on a medium supplemented with 2mM Mn^2+^ or on control plates without manganese. After ten days, the untreated seedlings showed a similar phenotype to the wild-type control Col-0 (Fig. 3c). *hflx1-1* mutant seedlings were slightly bigger than the wild type and other mutants, however this difference was not significant (Fig. 3d). Excess manganese caused a heterogenous response among all the lines tested. Some seedlings were only mildly affected, while others showed severe growth limitation, chlorophyll loss and cotyledon death (Fig 3c). No significant difference was observed between lines after 17 days of treatment (Fig 3d).

Next, we investigated the effect of cold and salt stress, two stresses known to perturb chloroplast gene expression (Zoschke & Bock, 2018; Hao et al., 2021). For cold stress, seedlings were transferred to 5°C and grown for 4 weeks (Fig. S3a). Growth was greatly inhibited during the first week and then resumed. The efficiency of photosystem II (Fv/Fm) showed a slight drop after cold stress treatment for all lines (Fig. S3b). However, no significant differences were detected between the mutants and the wild type. For salt stress treatment, seedlings were transferred to a medium supplemented with 150 mM NaCl. Four days after transfer, many seedlings became pale and showed evidence of cotyledon death. A similar phenotype was observed for both the wild type and the *hflx* mutants (Fig. S3c). Although *hflx1-1* seedlings showed a higher rate of cotyledon death, no significant difference was observed between lines (Fig. S3d). Overall, we found that HflX does not appear to be required for acclimation to heat, manganese, cold or salt stress.

### *hflx* mutants are sensitive to lincomycin

Next, we used lincomycin to directly inhibit chloroplast translation in seedlings. Strikingly, we found that the three *hflx* mutants were clearly more sensitive to lincomycin than the wild type (Fig. 4a). Lincomycin sensitivity was further confirmed by quantifying chlorophyll content which was significantly lower in the *hflx* mutants than in the wild-type control (Fig. 4b).

**Figure 4.**
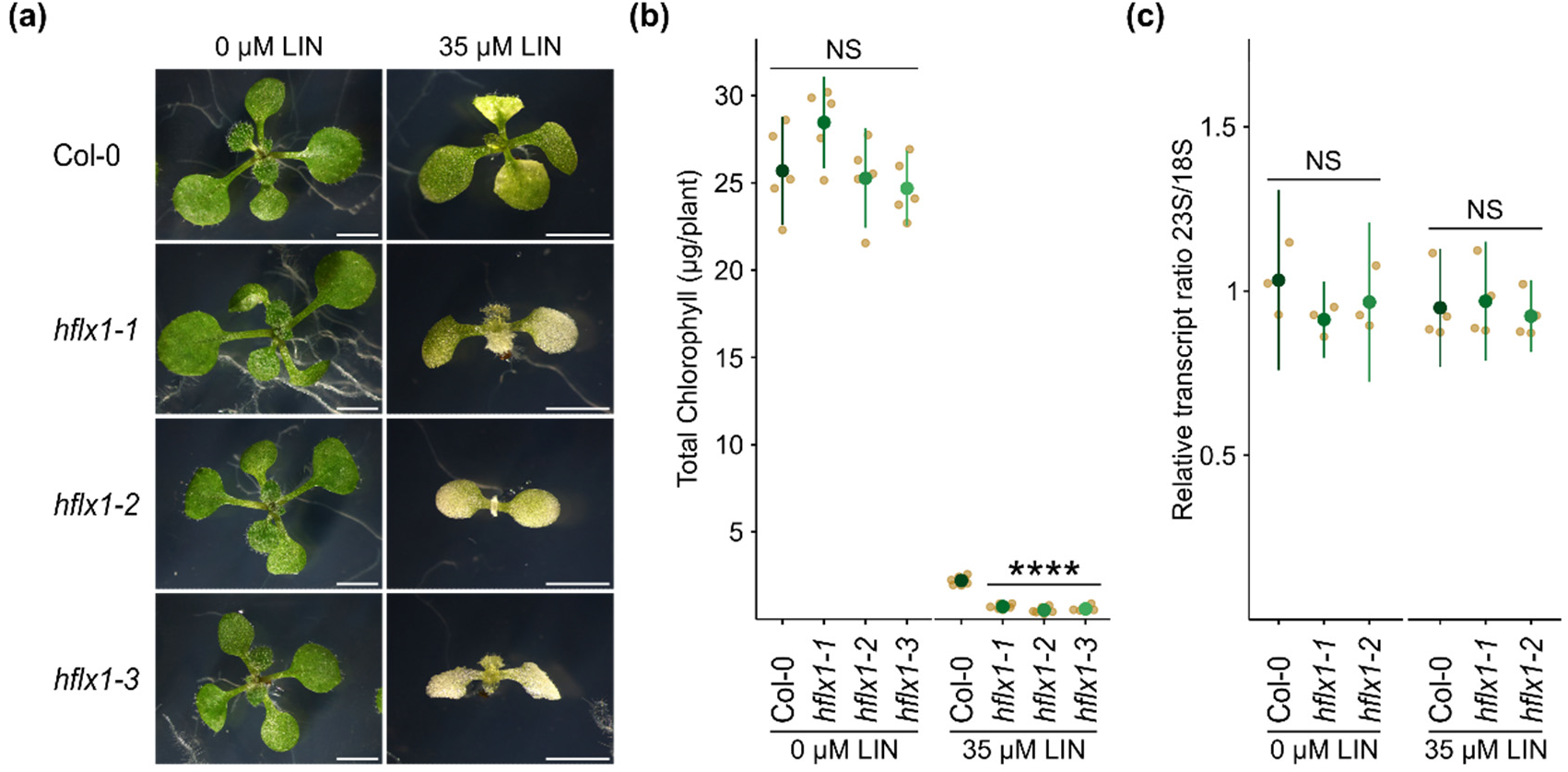
HflX is required for resistance to the antibiotic lincomycin. (a) Seedlings were grown on medium with or without lincomycin and photographed after 12 days. Scale bar, 3 mm. (b) Chlorophyll content was quantified in 15-day old seedlings, n = 5 biological replicates. (c) Ratio of the chloroplast 23S rRNA to the cytoplasmic 18S rRNA in Col-0, *hflx1-1* and *hflx1-2*. qRT-PCR was performed on cDNA extracted from seedlings grown on medium with or without lincomycin for 12 (untreated) or 15 (treated) days. n= 3-4 independent biological replicates. Graphs show mean and 95% CI. NS, not significant.

To determine whether the *hflx* lincomycin sensitivity was due to changes in chloroplast rRNA levels, we quantified the ratio of chloroplast 23S rRNA to cytosolic 18S rRNA by qPCR (Fig 4c). The 23S/18S rRNA ratio was similar between Col-0, *hflx1-1* and *hflx1-2* under both control and lincomycin treatment conditions, and showed no significant differences. Therefore, *hflx* does not appear to be required for the build-up of chloroplast rRNA levels under control or stress conditions.

## Discussion

Recently, it was reported that two HflX homologs, HflX and HflXr, are found in bacteria (Duval et al., 2018; Koller et al., 2022). Our analyses confirmed that the HflX of green plants and algae is closer to HflX than HflXr (Fig. 1A). This is in line with the wider distribution of HflX, and the specific association of HflXr with antibiotic resistance. In addition, our results reinforce previous findings suggesting that the plant HflX originated from green non-sulfur bacteria through lateral gene transfer (Suwastika et al., 2014). Interestingly, we also found a second clade of plant HflX-like enzymes that lacks the CTD that characterizes the majority of HflX enzymes in plants, bacteria, and animals (Fig. 1A, S1). The CTD, which does not interface directly with the ribosome, is reported to be less conserved compared to the other domains. Indeed, the CTD is also absent from the HflX of the archaeon *Sulfolobus solfataricus* (Wu et al., 2010).

We show that the canonical HflX is not required for plant growth and development under normal conditions (Fig. 2, S2). This is rather similar to the *E. coli* HflX that is dispensable for normal growth (Zhang et al., 2015). In contrast, GTPBP6, the human ortholog of HflX, is essential for cell survival and gene expression under physiological conditions (Lavdovskaia et al., 2020). This is likely due to the essential role of GTPBP6 in the assembly of mitochondrial ribosomes.

The *E. coli* HflX possess an ATP-dependent RNA helicase activity in the NTD that has an essential role in restoring heat damaged ribosomes (Dey et al., 2018). Although this domain is conserved in Arabidopsis HflX, with conservation of structure and essential residues for ATPase activity, Arabidopsis *hflx* mutants did not display compromised heat tolerance (Fig. 3). Similarly we were unable to detect increased sensitivity to excess manganese, another phenotype observed in *E. coli* HflX mutants (Kaur et al., 2014; Sengupta et al., 2018). This suggests that Arabidopsis HflX is not involved in chloroplast ribosome rescue or manganese homeostasis under the conditions tested, or that HflX plays a redundant role that can be replaced by other factors. One possibility is that HflX-like, which possesses the necessary domains, might also contribute to heat and manganese stress acclimation. It would therefore be interesting to test the chloroplast localization of HflX-like, and investigate the phenotype of *hflx* and *hflx-like* double mutants.

We also show that, in addition to heat and manganese stress, *hflx* mutants are not hypersensitive to cold or salt stress (Fig. S3). These conditions are known to affect chloroplast function, and in particular chloroplast translation (Zoschke & Bock, 2018; Hu et al., 2020; Hao et al., 2021). Indeed, some chloroplast translation factors are known to be required under such conditions (Li et al., 2018; Pulido et al., 2018). This would suggest that the removal of HflX does not perturb chloroplast ribosome biogenesis or translation enough to cause a detectable phenotype.

The hypersensitive phenotype we found in response to lincomycin treatment strongly implies that HflX is indeed ribosome-associated, and likely involved in the surveillance of chloroplast translation. However, even though HflX is not essential, it is still uncertain whether it participates in ribosome biogenesis. We did not observe a reduction in chloroplast rRNA levels in the *hflx* mutants. However, this does not exclude involvement in the later steps of ribosome biogenesis.

For example, the human GTPBP6 mutant does not show alterations in the steady-state levels of mitochondrial rRNAs or severe changes in the steady state levels of ribosomal proteins, yet still shows critical defects in mitochondrial large subunit assembly (mitoLSU) (Lavdovskaia et al., 2020). Lincomycin inhibits prokaryotic protein synthesis by interacting with the peptidyl transferase center (PTC). HflX and HflXr resistance to macrolide-lincosamide antibiotics is conferred by their ability to split and recycle stalled ribosomes (Duval et al., 2018; Rudra et al., 2020). HflXr also employs a second resistance mechanism. The cryo-EM structure of HflXr reveals that it binds analogously to *E. coli* HflX on the 50s subunit, with the N-loop of the NTD positioned deeper within the PTC. Upon binding to the ribosome, HflXr induces conformational changes in the PTC that are incompatible with antibiotic binding (Koller et al., 2022). A similar mechanism was more recently observed for HflX mediated resistance to chloramphenicol in *E. coli* suggesting that it is not limited to HflXr (Wu et al., 2022). In line with this, and considering the structural differences of the chloroplast ribosome compared to the bacterial ribosome (Yamaguchi & Subramanian, 2000, 2003; Manuell et al., 2007; Sharma et al., 2007), the antibiotic sensitivity of *hflx* mutants might be due to better access of lincomycin to its binding site in the absence of HflX, or loss of the capacity of HflX to recycle lincomycin stalled ribosomes. We note that, in either case, HflX-like is not able to prevent lincomycin sensitivity. This may be because HflX-like is localized in the mitochondria rather than the chloroplast (Rugen et al., 2019), or because the atypical structure of HflX-like alters or even prevents ribosome binding.

In conclusion, our data suggest that Arabidopsis HflX is a conserved HflX orthologue that is associated with the chloroplast ribosome and is likely to play a role in the surveillance of chloroplast translation. Even so, Arabidopsis HflX seems not to be actively involved during stress acclimation, suggesting that it is likely redundant with other plant factors, or plays an unknown role. Our results highlight the challenges of exploring translation regulation within the chloroplast, and further emphasize that while the functions of some ribosome-associated proteins are evolutionary conserved between organelles and bacteria, others are likely to be organism-dependent.

## Acknowledgements

We thank our colleaguesxg for managing the plant growth facilities. We acknowledge funding form the Erasmus+ International Credit Mobility for MM and funding from the Agence Nationale de la Recherche (ANR-22-CE20-0033, ANR-17-CE13-0005).

## Supplementary figures

**Figure S1.**
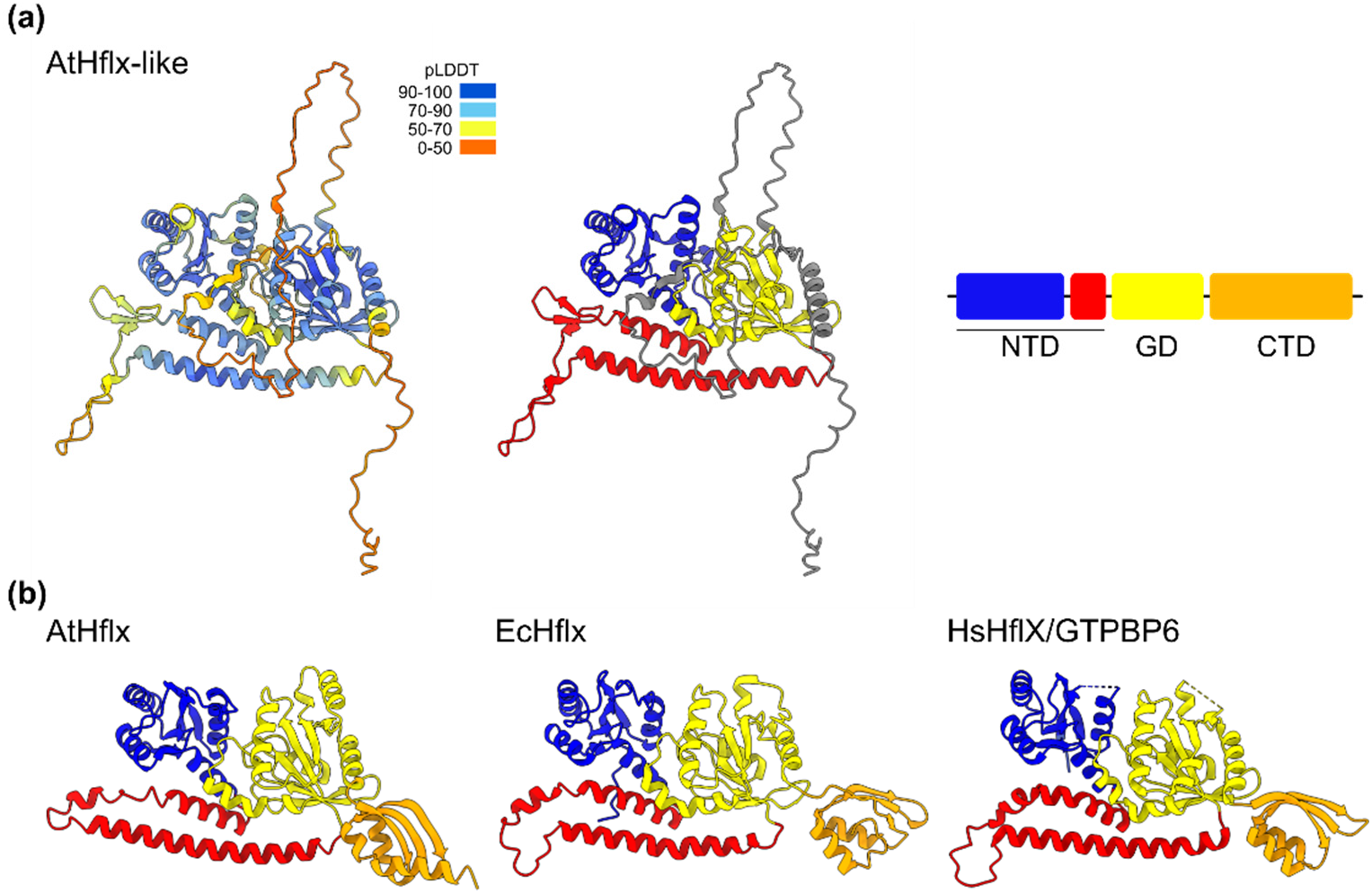
Structure of Arabidopsis HflX-like. (a) Alphafold model of Arabidopsis HflX-like (Q0WTB4) showing confidence-per-residue coloring (pLDDT) and domain organisation. The predicted chloroplast transit peptide is not shown. (b) The Alphafold model of the canonical Arabidopsis HflX (AtHflX, Q9FJM0), and the structures of ribosome-associated *E. coli* HflX (EcHflX, PDB 5ADY) and mitochondrial ribosome-associated *Homo sapiens* HflX/GTPB6 (HsHflX, PDB 7OF2)(Hillen et al., 2021). NTD, N-terminal domain; GD, G-domain; CTD, C-terminal domain.

**Figure S2.**
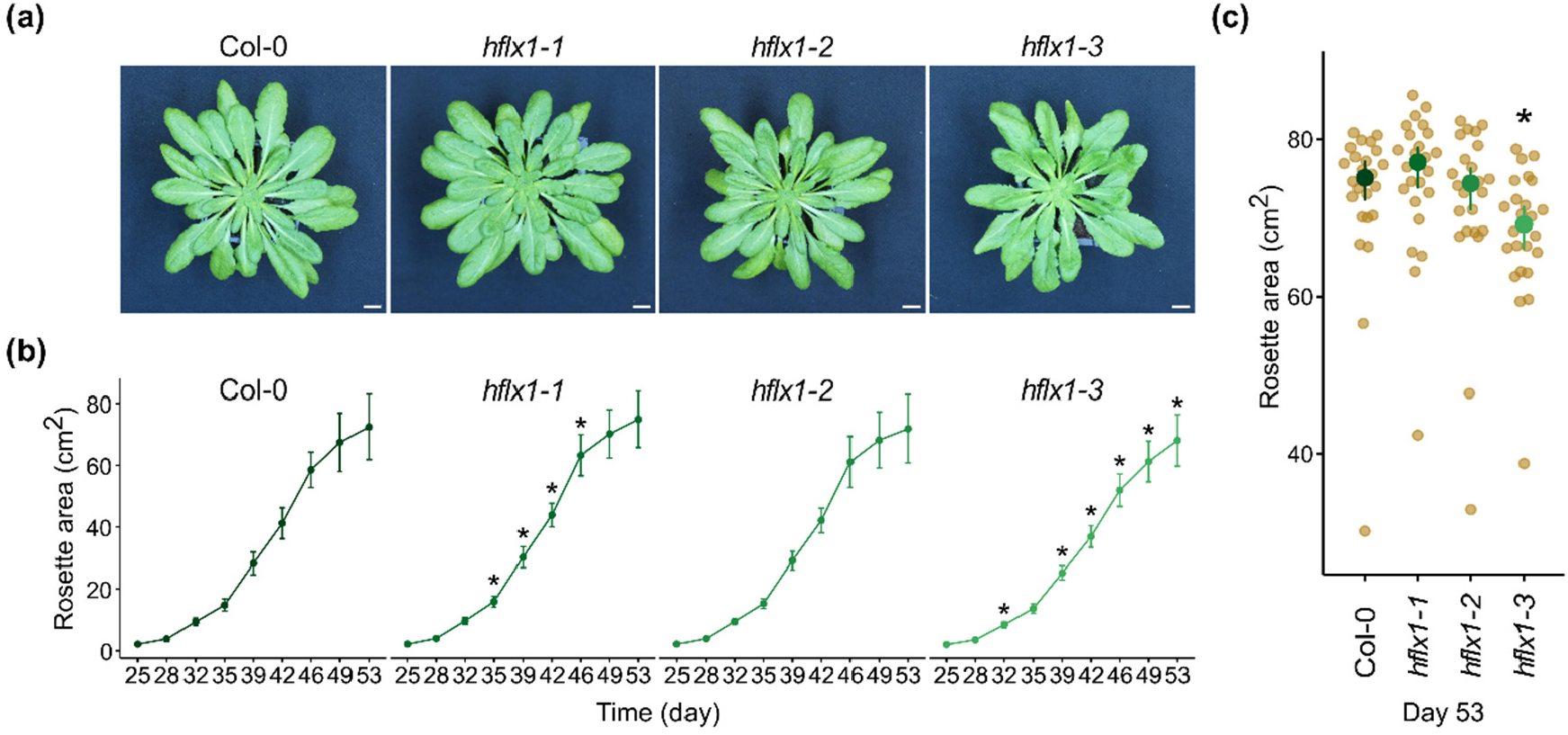
HflX is not required for vegetative growth under short days. The phenotype of wild type (Col-0) and *hflx* mutants grown in long day conditions. (a) Photographs of plant rosettes at day 53, (b) quantification of vegetative growth rates, and (c) comparison of rosette area at day 53 (n= 24 plants per genotype). Scale bar, 1cm. Graphs show mean and 95% CI. Statistical tests shown against Col-0.

**Fig. S3.**
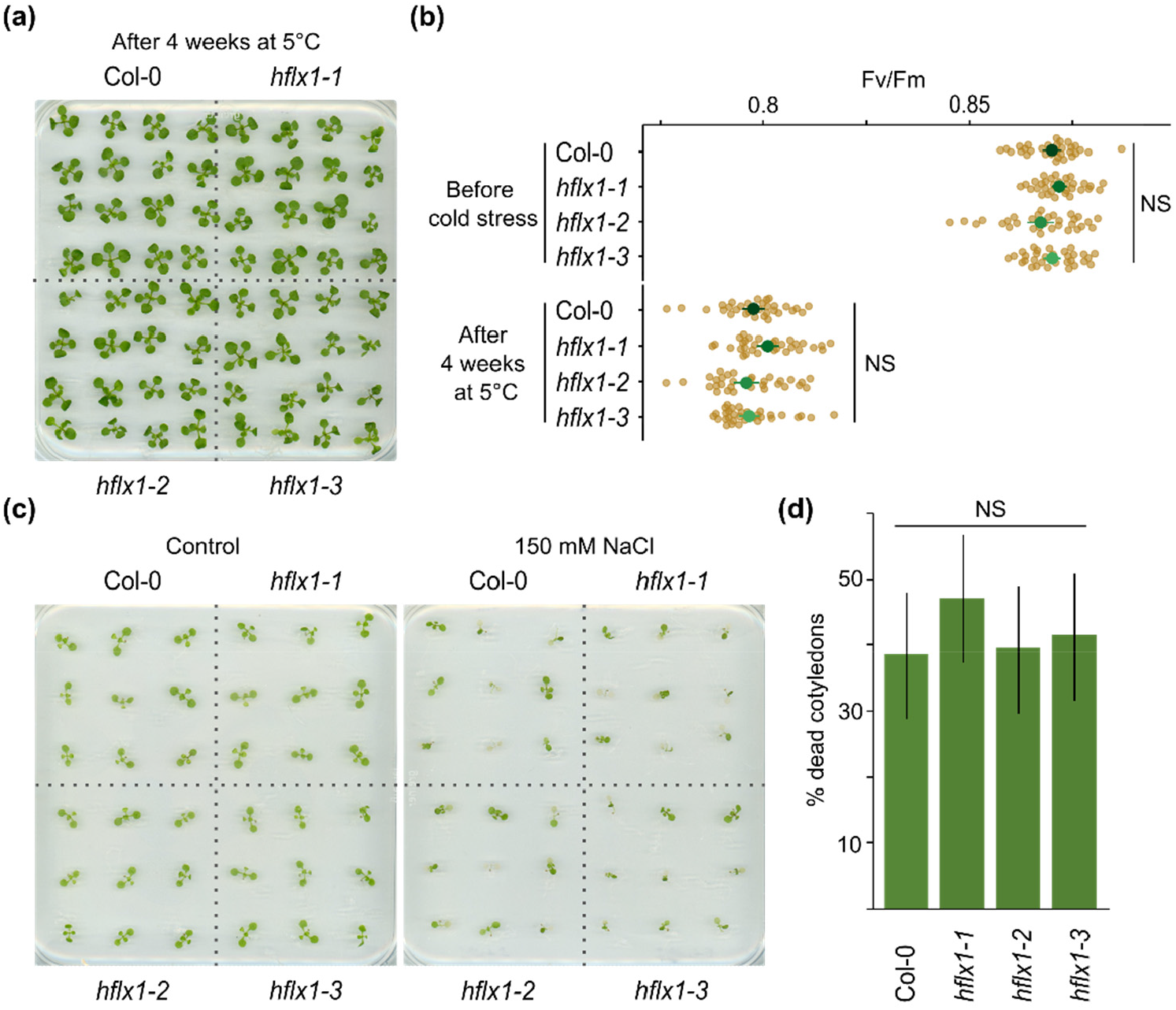
HflX is not required for acclimation to cold or salt stress. (a) One week-old seedlings were subjected to cold stress by transferring to 5°C for 4 weeks and photographed. (b) Fv/Fm in seedlings before and after cold stress treatment, n= 32 plants per genotype. (c) 8-day-old seedlings were transferred to media with or without 150 mM NaCl and photographed after 4 days. (d) Percentage of seedlings with at least one dead cotyledon from three pooled experimental replicates, n = 106 seedlings per genotype. Graphs show mean and 95% CI. Statistical tests shown against Col-0. NS, not significant.

**Table S1.**
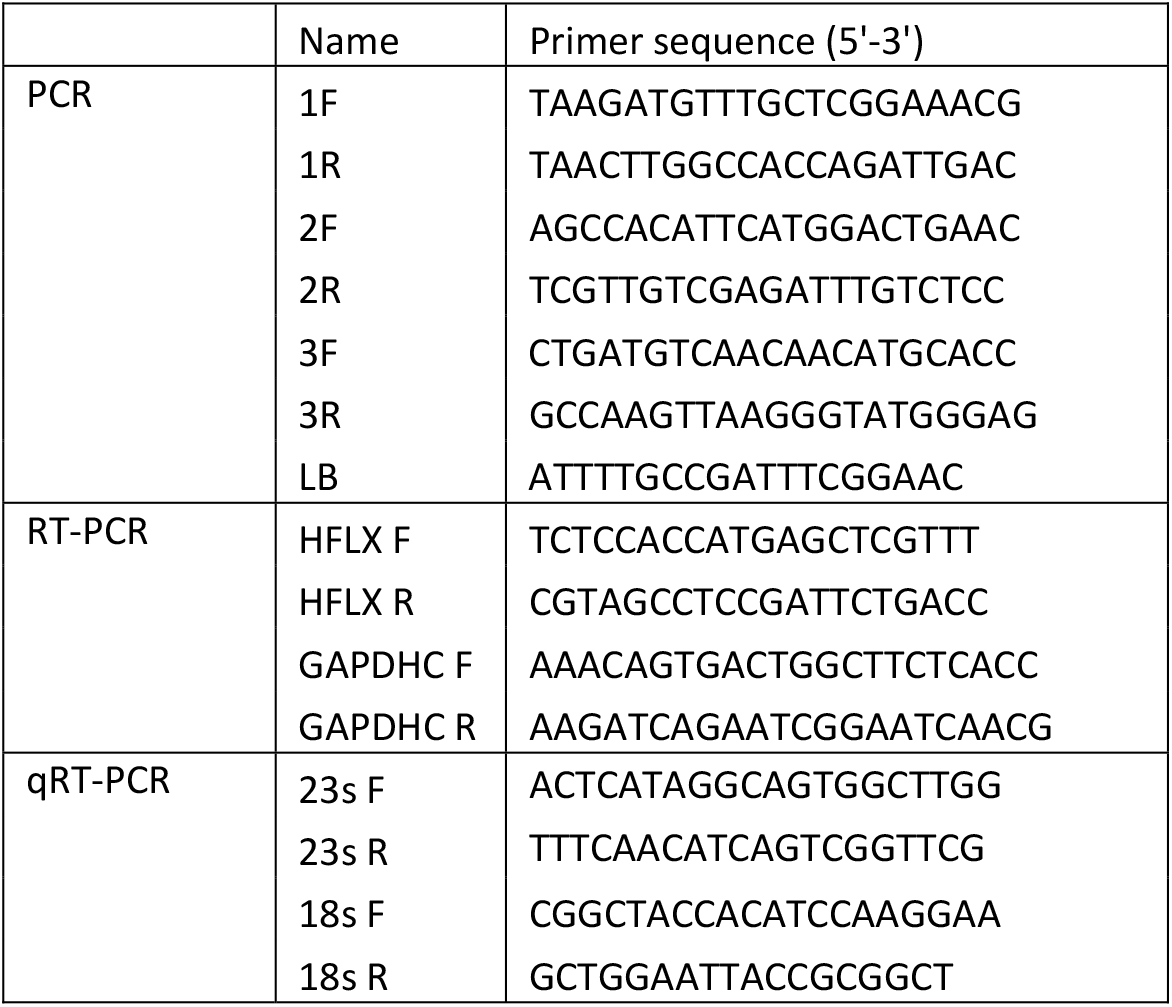

## Notes

### Competing Interest Statement

The authors have declared no competing interest.

## References

Almagro Armenteros, J.J., Salvatore, M., Emanuelsson, O., Winther, O., von Heijne, G., Elofsson, A., & Nielsen, H. (2019). Detecting sequence signals in targeting peptides using deep learning. Life Science Alliance, 2, e201900429. https://doi.org/10.26508/lsa.201900429

Alonso, J.M., Stepanova, A.N., Leisse, T.J., Kim, C.J., Chen, H., Shinn, P., Stevenson, D.K., Zimmerman, J., Barajas, P., Cheuk, R., Gadrinab, C., Heller, C., Jeske, A., Koesema, E., Meyers, C.C., Parker, H., Prednis, L., Ansari, Y., Choy, N., Deen, H., Geralt, M., Hazari, N., Hom, E., Karnes, M., Mulholland, C., Ndubaku, R., Schmidt, I., Guzman, P., Aguilar-Henonin, L., Schmid, M., Weigel, D., Carter, D.E., Marchand, T., Risseeuw, E., Brogden, D., Zeko, A., Crosby, W.L., Berry, C.C., & Ecker, J.R. (2003). Genome-wide insertional mutagenesis of Arabidopsis thaliana. Science (New York, N.Y.), 301, 653–657. https://doi.org/10.1126/science.1086391

Bang, W.Y., Chen, J., Jeong, I.S., Kim, S.W., Kim, C.W., Jung, H.S., Lee, K.H., Kweon, H.S., Yoko, I., Shiina, T., & Bahk, J.D. (2012). Functional characterization of ObgC in ribosome biogenesis during chloroplast development. Plant J., 71, 122–34. https://doi.org/10.1111/j.1365-313X.2012.04976.x

Chazaux, M., Schiphorst, C., Lazzari, G., & Caffarri, S. (2022). Precise estimation of chlorophyll a, b and carotenoid content by deconvolution of the absorption spectrum and new simultaneous equations for Chl determination. The Plant Journal, 109, 1630–1648. https://doi.org/10.1111/tpj.15643

Chigri, F., Sippel, C., Kolb, M., & Vothknecht, U.C. (2009). Arabidopsis OBG-Like GTPase (AtOBGL) Is Localized in Chloroplasts and Has an Essential Function in Embryo Development. Molecular Plant, 2, 1373–1383. https://doi.org/10.1093/mp/ssp073

Coatham, M.L., Brandon, H.E., Fischer, J.J., Schümmer, T., & Wieden, H.-J. (2016). The conserved GTPase HflX is a ribosome splitting factor that binds to the E-site of the bacterial ribosome. Nucleic Acids Research, 44, 1952–1961. https://doi.org/10.1093/nar/gkv1524

Dey, S., Biswas, C., & Sengupta, J. (2018). The universally conserved GTPase HflX is an RNA helicase that restores heat-damaged Escherichia coli ribosomes. Journal of Cell Biology, 217, 2519–2529. https://doi.org/10.1083/jcb.201711131

Duval, M., Dar, D., Carvalho, F., Rocha, E.P.C., Sorek, R., & Cossart, P. (2018). HflXr, a homolog of a ribosome-splitting factor, mediates antibiotic resistance. Proceedings of the National Academy of Sciences of the United States of America, 115, 13359–13364. https://doi.org/10.1073/pnas.1810555115

Eckardt, N.A., Snyder, G.W., Portis Jr, A.R., & Ogren, W.L. (1997). Growth and Photosynthesis under High and Low Irradiance of Arabidopsis thaliana Antisense Mutants with Reduced Ribulose-1,5-Bisphosphate Carboxylase/Oxygenase Activase Content. Plant Physiology, 113, 575–586. https://doi.org/10.1104/pp.113.2.575

Edwards, K., Johnstone, C., & Thompson, C. (1991). A simple and rapid method for the preparation of plant genomic DNA for PCR analysis.. Nucleic Acids Research, 19, 1349

Hao, S., Wang, Y., Yan, Y., Liu, Y., Wang, J., & Chen, S. (2021). A Review on Plant Responses to Salt Stress and Their Mechanisms of Salt Resistance. Horticulturae, 7, 132. https://doi.org/10.3390/horticulturae7060132

Hillen, H.S., Lavdovskaia, E., Nadler, F., Hanitsch, E., Linden, A., Bohnsack, K.E., Urlaub, H., & Richter-Dennerlein, R. (2021). Structural basis of GTPase-mediated mitochondrial ribosome biogenesis and recycling. Nature Communications, 12, 3672. https://doi.org/10.1038/s41467-021-23702-y

Hu, S., Ding, Y., & Zhu, C. (2020). Sensitivity and Responses of Chloroplasts to Heat Stress in Plants. Frontiers in Plant Science, 11

Hüther, P., Schandry, N., Jandrasits, K., Bezrukov, I., & Becker, C. (2020). ARADEEPOPSIS, an Automated Workflow for Top-View Plant Phenomics using Semantic Segmentation of Leaf States. The Plant Cell, 32, 3674–3688. https://doi.org/10.1105/tpc.20.00318

Jain, N., Dhimole, N., Khan, A.R., De, D., Tomar, S.K., Sajish, M., Dutta, D., Parrack, P., & Prakash, B. (2009). E. coli HflX interacts with 50S ribosomal subunits in presence of nucleotides. Biochemical and Biophysical Research Communications, 379, 201–205. https://doi.org/10.1016/j.bbrc.2008.12.072

Jumper, J., Evans, R., Pritzel, A., Green, T., Figurnov, M., Ronneberger, O., Tunyasuvunakool, K., Bates, R., Žídek, A., Potapenko, A., Bridgland, A., Meyer, C., Kohl, S.A.A., Ballard, A.J., Cowie, A., Romera-Paredes, B., Nikolov, S., Jain, R., Adler, J., Back, T., Petersen, S., Reiman, D., Clancy, E., Zielinski, M., Steinegger, M., Pacholska, M., Berghammer, T., Bodenstein, S., Silver, D., Vinyals, O., Senior, A.W., Kavukcuoglu, K., Kohli, P., & Hassabis, D. (2021). Highly accurate protein structure prediction with AlphaFold. Nature, 596, 583–589. https://doi.org/10.1038/s41586-021-03819-2

Katoh, K., Rozewicki, J., & Yamada, K.D. (2019). MAFFT online service: multiple sequence alignment, interactive sequence choice and visualization. Briefings in Bioinformatics, 20, 1160–1166. https://doi.org/10.1093/bib/bbx108

Kaur, G., Sengupta, S., Kumar, V., Kumari, A., Ghosh, A., Parrack, P., & Dutta, D. (2014). Novel MntR-independent mechanism of manganese homeostasis in Escherichia coli by the ribosome-associated protein HflX. Journal of Bacteriology, 196, 2587–2597. https://doi.org/10.1128/JB.01717-14

Koller, T.O., Turnbull, K.J., Vaitkevicius, K., Crowe-McAuliffe, C., Roghanian, M., Bulvas, O., Nakamoto, J.A., Kurata, T., Julius, C., Atkinson, G.C., Johansson, J., Hauryliuk, V., & Wilson, D.N. (2022). Structural basis for HflXr-mediated antibiotic resistance in Listeria monocytogenes. Nucleic Acids Research, 50, 11285–11300. https://doi.org/10.1093/nar/gkac934

Lavdovskaia, E., Denks, K., Nadler, F., Steube, E., Linden, A., Urlaub, H., Rodnina, M.V., & Richter-Dennerlein, R. (2020). Dual function of GTPBP6 in biogenesis and recycling of human mitochondrial ribosomes. Nucleic Acids Research, 48, 12929–12942. https://doi.org/10.1093/nar/gkaa1132

Li, X., Cai, C., Wang, Z., Fan, B., Zhu, C., & Chen, Z. (2018). Plastid Translation Elongation Factor Tu Is Prone to Heat-Induced Aggregation Despite Its Critical Role in Plant Heat Tolerance. Plant Physiology, 176, 3027–3045. https://doi.org/10.1104/pp.17.01672

Liu, S., Zheng, L., Jia, J., Guo, J., Zheng, M., Zhao, J., Shao, J., Liu, X., An, L., Yu, F., & Qi, Y. (2019). Chloroplast Translation Elongation Factor EF-Tu/SVR11 Is Involved in var2-Mediated Leaf Variegation and Leaf Development in Arabidopsis. Frontiers in Plant Science, 10

Makino, A., & Osmond, B. (1991). Effects of Nitrogen Nutrition on Nitrogen Partitioning between Chloroplasts and Mitochondria in Pea and Wheat 1. Plant Physiology, 96, 355–362. https://doi.org/10.1104/pp.96.2.355

Manuell, A.L., Quispe, J., & Mayfield, S.P. (2007). Structure of the Chloroplast Ribosome: Novel Domains for Translation Regulation. PLOS Biology, 5, e209. https://doi.org/10.1371/journal.pbio.0050209

Mehrez, M., Romand, S., & Field, B. (2023). New perspectives on the molecular mechanisms of stress signalling by the nucleotide guanosine tetraphosphate (ppGpp), an emerging regulator of photosynthesis in plants and algae. New Phytologist, 237, 1086–1099. https://doi.org/10.1111/nph.18604

Olinares, P.D.B., Ponnala, L., & van Wijk, K.J. (2010). Megadalton Complexes in the Chloroplast Stroma of Arabidopsis thaliana Characterized by Size Exclusion Chromatography, Mass Spectrometry, and Hierarchical Clustering*. Molecular & Cellular Proteomics, 9, 1594–1615. https://doi.org/10.1074/mcp.M000038-MCP201

Pettersen, E.F., Goddard, T.D., Huang, C.C., Meng, E.C., Couch, G.S., Croll, T.I., Morris, J.H., & Ferrin, T.E. (2021). UCSF ChimeraX: Structure visualization for researchers, educators, and developers. Protein Science : A Publication of the Protein Society, 30, 70–82. https://doi.org/10.1002/pro.3943

Pulido, P., Zagari, N., Manavski, N., Gawronski, P., Matthes, A., Scharff, L.B., Meurer, J., & Leister, D. (2018). CHLOROPLAST RIBOSOME ASSOCIATED Supports Translation under Stress and Interacts with the Ribosomal 30S Subunit. Plant Physiology, 177, 1539–1554. https://doi.org/10.1104/pp.18.00602

Romand, S., Abdelkefi, H., Lecampion, C., Belaroussi, M., Dussenne, M., Ksas, B., Citerne, S., Caius, J., D’Alessandro, S., Fakhfakh, H., Caffarri, S., Havaux, M., & Field, B. (2022). A guanosine tetraphosphate (ppGpp) mediated brake on photosynthesis is required for acclimation to nitrogen limitation in Arabidopsis. eLife, 11, e75041. https://doi.org/10.7554/eLife.75041

Rudra, P., Hurst-Hess, K.R., Cotten, K.L., Partida-Miranda, A., & Ghosh, P. (2020). Mycobacterial HflX is a ribosome splitting factor that mediates antibiotic resistance. Proceedings of the National Academy of Sciences, 117, 629–634. https://doi.org/10.1073/pnas.1906748117

Rugen, N., Straube, H., Franken, L.E., Braun, H.-P., & Eubel, H. (2019). Complexome Profiling Reveals Association of PPR Proteins with Ribosomes in the Mitochondria of Plants. Molecular & Cellular Proteomics : MCP, 18, 1345–1362. https://doi.org/10.1074/mcp.RA119.001396

Schaefer, L., Uicker, W.C., Wicker-Planquart, C., Foucher, A.-E., Jault, J.-M., & Britton, R.A. (2006). Multiple GTPases Participate in the Assembly of the Large Ribosomal Subunit in Bacillus subtilis. Journal of Bacteriology, 188, 8252–8258. https://doi.org/10.1128/JB.01213-06

Sengupta, S., Mondal, A., Dutta, D., & Parrack, P. (2018). HflX protein protects Escherichia coli from manganese stress. Journal of Biosciences, 43, 1001–1013

Sharma, M.R., Wilson, D.N., Datta, P.P., Barat, C., Schluenzen, F., Fucini, P., & Agrawal, R.K. (2007). Cryo-EM study of the spinach chloroplast ribosome reveals the structural and functional roles of plastid-specific ribosomal proteins. Proceedings of the National Academy of Sciences, 104, 19315–19320. https://doi.org/10.1073/pnas.0709856104

Sugliani, M., Abdelkefi, H., Ke, H., Bouveret, E., Robaglia, C., Caffarri, S., & Field, B. (2016). An Ancient Bacterial Signaling Pathway Regulates Chloroplast Function to Influence Growth and Development in Arabidopsis. The Plant Cell, 28, 661–679. https://doi.org/10.1105/tpc.16.00045

Suwastika, I.N., Denawa, M., Yomogihara, S., Im, C.H., Bang, W.Y., Ohniwa, R.L., Bahk, J.D., Takeyasu, K., & Shiina, T. (2014). Evidence for lateral gene transfer (LGT) in the evolution of eubacteria-derived small GTPases in plant organelles. Frontiers in Plant Science, 5

Trifinopoulos, J., Nguyen, L.-T., von Haeseler, A., & Minh, B.Q. (2016). W-IQ-TREE: a fast online phylogenetic tool for maximum likelihood analysis. Nucleic Acids Research, 44, W232–W235. https://doi.org/10.1093/nar/gkw256

Varadi, M., Anyango, S., Deshpande, M., Nair, S., Natassia, C., Yordanova, G., Yuan, D., Stroe, O., Wood, G., Laydon, A., Žídek, A., Green, T., Tunyasuvunakool, K., Petersen, S., Jumper, J., Clancy, E., Green, R., Vora, A., Lutfi, M., Figurnov, M., Cowie, A., Hobbs, N., Kohli, P., Kleywegt, G., Birney, E., Hassabis, D., & Velankar, S. (2022). AlphaFold Protein Structure Database: massively expanding the structural coverage of protein-sequence space with high-accuracy models. Nucleic Acids Research, 50, D439–D444. https://doi.org/10.1093/nar/gkab1061

Wu, D., Dai, Y., & Gao, N. (2022). Cryo-EM Structure of the 50S-HflX Complex Reveals a Novel Mechanism of Antibiotic Resistance in E. coli 2022.11.25.517942. https://doi.org/10.1101/2022.11.25.517942

Wu, H., Sun, L., Blombach, F., Brouns, S.J.J., Snijders, A.P.L., Lorenzen, K., van den Heuvel, R.H.H., Heck, A.J.R., Fu, S., Li, X., Zhang, X.C., Rao, Z., & van der Oost, J. (2010). Structure of the ribosome associating GTPase HflX. Proteins, 78, 705–713. https://doi.org/10.1002/prot.22599

Yamaguchi, K., & Subramanian, A.R. (2000). The Plastid Ribosomal Proteins: IDENTIFICATION OF ALL THE PROTEINS IN THE 50 S SUBUNIT OF AN ORGANELLE RIBOSOME (CHLOROPLAST) *. Journal of Biological Chemistry, 275, 28466–28482. https://doi.org/10.1074/jbc.M005012200

Yamaguchi, K., & Subramanian, A.R. (2003). Proteomic identification of all plastid-specific ribosomal proteins in higher plant chloroplast 30S ribosomal subunit. European Journal of Biochemistry, 270, 190–205. https://doi.org/10.1046/j.1432-1033.2003.03359.x

Zhang, Y., Mandava, C.S., Cao, W., Li, X., Zhang, D., Li, N., Zhang, Y., Zhang, X., Qin, Y., Mi, K., Lei, J., Sanyal, S., & Gao, N. (2015). HflX is a ribosome-splitting factor rescuing stalled ribosomes under stress conditions. Nature Structural & Molecular Biology, 22, 906–913. https://doi.org/10.1038/nsmb.3103

Zoschke, R., & Bock, R. (2018). Chloroplast Translation: Structural and Functional Organization, Operational Control, and Regulation. The Plant Cell, 30, 745–770. https://doi.org/10.1105/tpc.18.00016

