## Supplementary Figures and Tables for "Insights into the function of the chloroplastic ribosome-associated GTPase HflX in *Arabidopsis thaliana*"

### Supplementary Tables and Figures

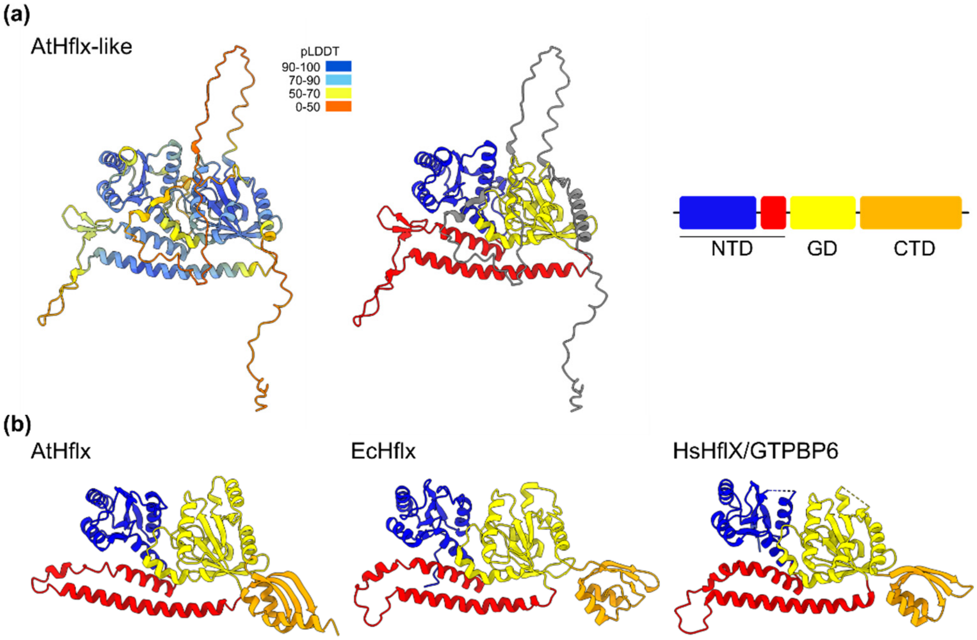


**Figure S1. Structure of Arabidopsis HflX-like.** (a) Alphafold model of Arabidopsis HflX-like (Q0WTB4) showing confidence-per-residue coloring (pLDDT) and domain organisation. The predicted chloroplast transit peptide is not shown. (b) The Alphafold model of the canonical Arabidopsis HflX (AtHflX, Q9FJM0), and the structures of ribosome-associated *E. coli* HflX (EcHflX, PDB 5ADY) and mitochondrial ribosome-associated *Homo sapiens* HflX/GTPB6 (HsHflX, PDB 7OF2)(Hillen et al., 2021). NTD, N-terminal domain; GD, G-domain; CTD, C-terminal domain.

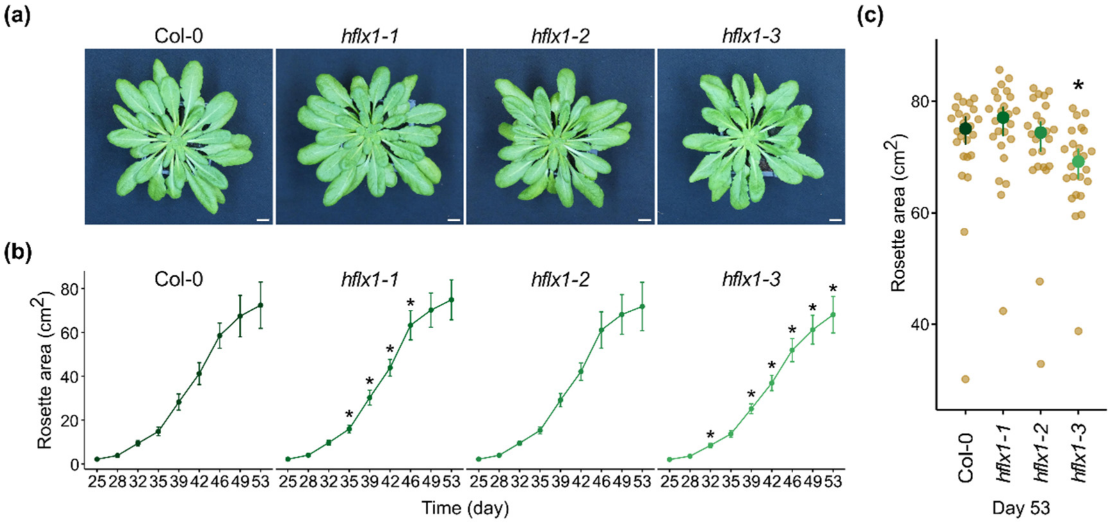


**Figure S2. HflX is not required for vegetative growth under short days.** The phenotype of wild type (Col-0) and *hflx* mutants grown in long day conditions. (a) Photographs of plant rosettes at day 53, (b) quantification of vegetative growth rates, and (c) comparison of rosette area at day 53 (n= 24 plants per genotype). Scale bar, 1cm. Graphs show mean and 95% CI. Statistical tests shown against Col-0.

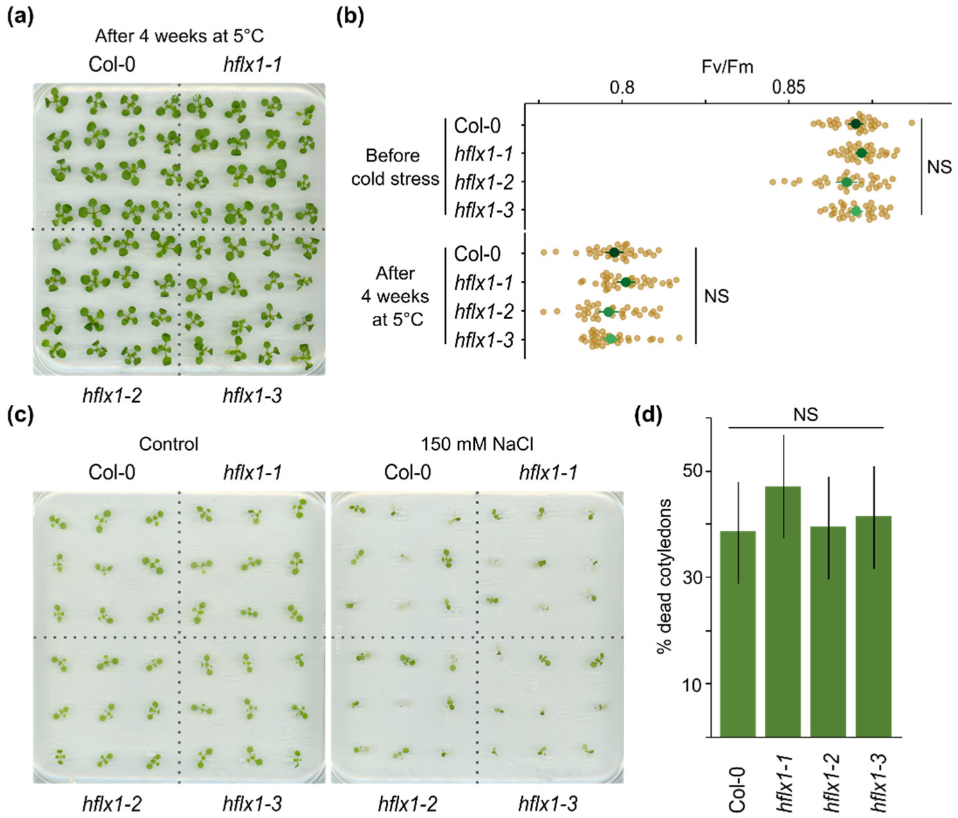


**Fig. S3. HflX is not required for acclimation to cold or salt stress.** (a) One week-old seedlings were subjected to cold stress by transferring to 5°C for 4 weeks and photographed. (b) Fv/Fm in seedlings before and after cold stress treatment, n= 32 plants per genotype. (c) 8-day-old seedlings were transferred to media with or without 150 mM NaCl and photographed after 4 days. (d) Percentage of seedlings with at least one dead cotyledon from three pooled experimental replicates, n = 106 seedlings per genotype. Graphs show mean and 95% CI. Statistical tests shown against Col-0. NS, not significant.

| | Name | Primer sequence (5'-3') |
|---|---|---|
| PCR | 1F | TAAGATGTTTGCTCGGAAACG |
| | 1R | TAACTTGGCCACCAGATTGAC |
| | 2F | AGCCACATTCATGGACTGAAC |
| | 2R | TCGTTGTCGAGATTTGTCTCC |
| | 3F | CTGATGTCAACAACATGCACC |
| | 3R | GCCAAGTTAAGGGTATGGGAG |
| | LB | ATTTTGCCGATTTCGGAAC |
| RT-PCR | HFLX F | TCTCCACCATGAGCTCGTTT |
| | HFLX R | CGTAGCCTCCGATTCTGACC |
| | GAPDHC F | AAACAGTGACTGGCTTCTCACC |
| | GAPDHC R | AAGATCAGAATCGGAATCAACG |
| qRT-PCR | 23s F | ACTCATAGGCAGTGGCTTGG |
| | 23s R | TTTCAACATCAGTCGGTTCG |
| | 18s F | CGGCTACCACATCCAAGGAA |
| | 18s R | GCTGGAATTACCGCGGCT |

**Table S1**
